# Proteome-wide Prediction of Lysine Methylation Reveals Novel Histone Marks and Outlines the Methyllysine Proteome

**DOI:** 10.1101/274688

**Authors:** Kyle K Biggar, Francois Charih, Huadong Liu, Yasser B Ruiz-Blanco, Leanne Stalker, Anand Chopra, Justin Connolly, Hemanta Adhikary, Kristin Frensemier, Marek Galka, Qi Fang, Christopher Wynder, William L Standford, James R Green, Shawn S-C. Li

**Author notes:** These authors contributed equally to the work (Kyle K Biggar, Francois Charih).

## Abstract

Protein Lys methylation plays a critical role in numerous cellular processes, yet it has been challenging to identify Lys methylation in a systematic manner. We present here an approach combining *in silico* prediction with targeted mass spectrometry (MS) to identify Lys methylation (Kme) sites at the proteome level. We have developed MethylSight, a program that predicts Kme events solely on physicochemical and biochemical properties of putative methylation sites, which can then be validated by targeted MS. Using this approach, we have identified 70 new histone Kme marks with a 90% validation rate. H2BK43me2, which undergoes dynamic changes during stem cell differentiation, is found to be a substrate of KDM5b. Furthermore, MethylSight predicts ~50,000 Kme sites in non-histone proteins with high confidence, suggesting that Lys methylation is a prevalent post-translational modification. Our work provides a useful resource for systematic exploration of the role of Lys methylation in human health and disease.

## INTRODUCTION

Post-translational modifications (PTMs) are reversible biochemical modifications that play a crucial role in the regulation of protein function and the transmission of biological signals (Mann and Jensen, 2003). To date, numerous different PTMs have been identified in human proteins, greatly expanding the size of the functional human proteome and endowing cells with the ability to regulate complex and dynamic cellular processes. Lysine (Lys) methylation is one such prevalent PTM that has been well established to occur on histones, but in recent years has been documented to occur on an expanding catalogue of non-histone proteins (Biggar and Li, 2015). The N-terminal tails of the core histone proteins are subject to methylation on numerous Lys residues; and together with other PTMs they comprise what is commonly referred to as the “histone-code” (Jones, 2016) that regulates chromatin structure and gene expression programs (Martin and Zhang, 2005). Although Lys methylation is commonly found on histones, non-histone protein methylation has received considerable attention in recent years due to its prevalence and role in regulating cellular functions under physiological and pathological conditions, including cancer (Biggar and Li, 2015; Guo et al., 2019; Hamamoto et al., 2015; Li et al., 2018; Reynoird et al., 2016; Yoshioka et al., 2016). Given the prevalence of Lys methylation and the technical challenges of identifying this PTM (Cao and Garcia, 2016; Cao et al., 2013; Carlson et al., 2014; Lund et al., 2019; Wang et al., 2018), it is of great interest to develop proteomic and computational approaches that allow for the systematic identification of Lys methylation. Along this line, several recent studies, employing *in silico* and mass spectrometry-based methods, have shed important light on the methyl-Lys proteome (Audagnotto and Dal Peraro, 2017; Cao and Garcia, 2016; Cao et al., 2013; Carlson et al., 2014; Liu et al., 2013; Lund et al., 2019; Shi et al., 2015a; Wang et al., 2018; Wen et al., 2016).

The number of known methylated proteins and modification sites has grown tremendously in recent years. Indeed, recent advances in identification technologies, including affinity enrichment methods and high-resolution mass spectrometry, have provided insight into non-histone protein Lys methylation, with many of these methylation events shown to have important regulatory functions for the respective proteins (Liu et al., 2013; Mazur et al., 2014; Thandapani et al., 2017). Furthermore, it is now known that dynamic protein methylation is involved in a growing number of cellular processes (Biggar and Li, 2015; Cornett et al., 2019; Wu et al., 2017). For example, the tumor suppressor p53 is methylated on multiple Lys residues and the dynamics and combinations of these modifications have the capacity to regulate p53 function through a surprisingly diverse array of mechanisms (West and Gozani, 2011). Furthermore, the methylation status of the catalytic subunit of DNA-dependent protein kinase (DNA-PKcs), an important regulator of DNA damage repair, has been shown to dictate. its ability to effectively repair damaged DNA (Liu et al., 2013). Together, these studies suggest a broad role for Lys methylation in regulating protein function, well beyond controlling chromatin dynamics via histone methylation.

The emergence of Lys methylation as an important regulator of cellular function calls for the development of approaches for the systematic identification of this PTM. One of the greatest challenges in the discovery of Lys-methylated proteins is the isolation and enrichment of the methylated proteins or peptides prior to their identification. In this regard, tt has proven difficult to develop specific affinity reagents for Lys methylation (Liu et al., 2013; Carlson et al., 2014; Wang et al., 2018). As a result, the identification of new Lys methylation sites has not experienced the same explosive growth in discovery as some other PTMs, such as serine/threonine and tyrosine phosphorylation, Lys acetylation, and arginine methylation. Although several affinity strategies that utilize natural methyl-binding domains have been remarkably successful in identifying new Lys methylation events when coupled with mass spectrometry (Liu et al., 2013; Carlson et al., 2014), these approaches are inherently biased towards the binding specificity of the domain used in the initial enrichment. *In silico* prediction methods overcome this limitation by uncovering methylation events solely based on the general underlying biochemical characteristics of known modification sites. We therefore set out to develop an integrated platform combining *in silico* prediction with targeted mass spectrometry (MS) to identify new Lys methylation events and expand the potential landscape of the methyl-Lys proteome.

During the past decade, there have been several attempts to develop methyl-Lys and methyl-arginine computational predictors (Table S1) (Chen et al., 2006; Hu et al., 2011; Qiu et al., 2014; Shao et al., 2009; Shi et al., 2012a; Shien et al., 2009, Deng et al., 2017, Lee et al., 2014). These studies built their models on a few hundred methylation sites extracted from the UniProtKB, PhosphoSitePlus, or/and PubMed databases. Based on the available information on protein methylation at the time of the study, the predictors were limited to approximately 200 non-redundant methyl-Lys sites for building and assessing the respective models. Given the rapid growth of validated methylated sites in recent years (Cao and Garcia, 2016) and the thousands of additional sites predicted to exist in human cells, it is unlikely for the models built on a few hundred known sites to capture the diversity of methyl-Lys sites. More importantly, methylation sites predicted from such *in silico* methods have, in essentially all cases, not been validated experimentally. Thus their potential in the identification of new cellular Lys methylation events has not been vigorously tested.

Through the use of a unique machine learning approach to predict new Lys methylation sites, we have developed a method that overcomes the limitations mentioned above. This is achieved partially by utilizing sequence alignment-free features (or descriptors) to directly capture the physical and chemical properties of the methylated sites, rather than structure-specific features that often fail due to the limited amount of available data. Moreover, the training dataset employed in our study contains more than three thousand sites collected from the PhosphoSitePlus database (http://phosphosite.org). Furthermore, we treat imbalance using cost-sensitive learning. Rather than introducing synthetic training data or losing valuable exemplars through under-sampling, the datasets were kept in their intrinsic imbalanced ratio during cross-validation and hold-out tests.

We show that the resulting method, MethylSight, can predict novel Lys methylation (Kme) sites in histone and non-histone proteins with high confidence. The predicted Kme sites can then be validated in cells or tissues by targeted MS such as Multiple Reaction Monitoring (MRM) or Parallel Reaction Monitoring (PRM) (Fig. 1) (Liu et al., 2013). We generated the first comprehensive map of methyl-Lys proteome, including both known and novel sites predicted to exist by MethylSight with high confidence. Our work suggests that the methyl-Lys proteome is as large as the phospho-tyrosine proteome. Besides revealing novel histone marks in histones H1 and H2, we have identified ~50,000 high-probability Kme sites in non-histone proteins involved in a diverse array of cellular processes and molecular functions. To enable the rapid growth of methyl-Lys research and the discovery of new functional methylation events, the MethylSight web server and datasets are made freely accessible via http://methylsight.com.

**Figure 1.**
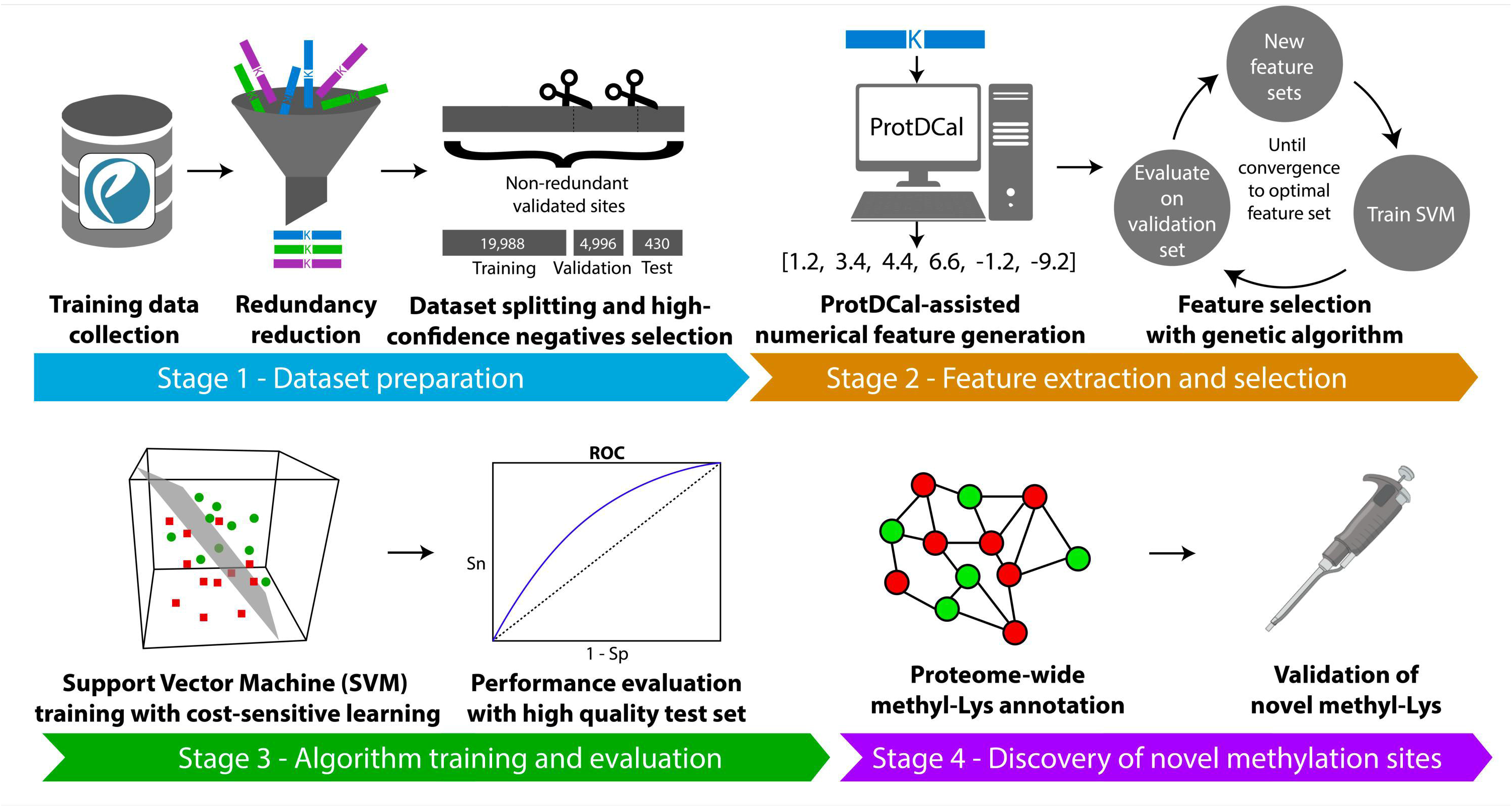
An overview of MethylSight. MethylSight enables proteome-wide prediction and systematic annotation of Lys methylation through four steps. First, published Lys methylation data are collected and selected to optimize training and testing. Second, numerical features describing potential methylation sites are generated, and the optimal subset of features is selected to optimize performance on a validation set. Third, a support vector machine model is trained with a cost-sensitive learning strategy and tested using low redundancy training and test sets. Finally, the methyl-Lys proteome is annotated to guide experimental validation of new methylation events.

## RESULTS

### The MethySight pipeline for global Lys methylation analysis

The MethySight prediction and validation pipeline comprised four main stages (Fig. 1). We started with the Lys methylation (Kme) site database collected at the PhosphoSitePlus (accessed 09/28/18) (Hornbeck et al., 2015). Upon removing redundant Kme sites, the positive Kme sites/peptides were divided randomly into training set, validation set, and test set. Lys residues within a methylated protein, but which are not reported to be methylated, were used as negatives in our analysis (Table 1; see Methods for details of selection for positives and negatives). Next, we used ProtDCal (Ruiz-Blanco et al., 2015, Romero-Molina et al., 2019) to generate a feature vector for each Lys-centered window in the dataset. The feature vector consists of a set of scalar values computed from the physico-chemical characteristics of amino acids using a variety of aggregation operations to capture the properties of a given amino acid and its neighborhood. At Stage 3, we employed support vector machine (SVM) models to predict how likely a Lys is to be methylated, based on the most relevant ProtDCal descriptors for a given Lys-centered window. The cost-sensitive sequential minimal optimization (SMO) algorithm (Cai and Cherkassky, 2012) was used to train all SVM classifiers in this work. Following a series of supervised and unsupervised steps to determine the optimal feature set (described in the Methods Details), we performed a grid search to optimize the SVM model hyperparameters. The hyperparameters were varied iteratively, and the set of hyperparameters producing the best prediction accuracy in 5-fold cross-validation over the training set, as well as on the validation set (see Table 1), were selected.

**Table 1.**
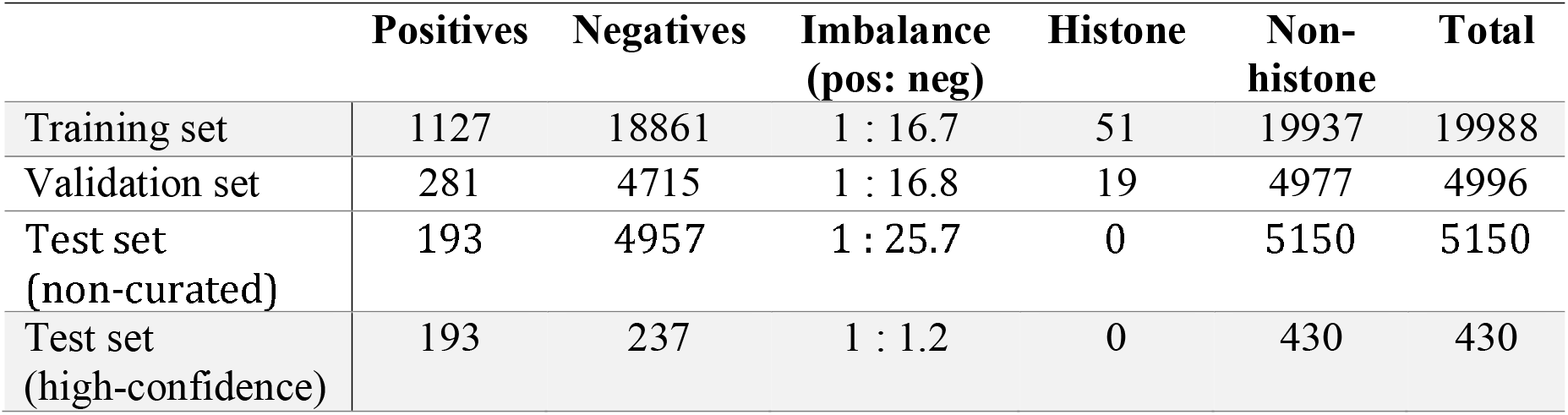
Composition of the datasets used to train, validate and test MethylSight.

As with previous studies (Chen et al., 2006; Deng et al., 2017; Hu et al., 2011; Lee et al., 2014; Qiu et al., 2014; Shao et al., 2009; Shi et al., 2012a; Shien et al., 2009), the non-methylated Lys residues in a methylated protein are here assumed to be negative when training and evaluating predictors. Considering that the number of methylation sites continues to grow rapidly, this assumption may be flawed, as many of the assumed-negative instances may actually be undocumented positive sites. This leads to a pessimistic estimation of the precision of the obtained model. Therefore, we also computed the precision using a high confidence negative test subset (see Methods Details). Shown in Fig. 2A, this can be considered an optimistic estimator of prediction precision, assuming that roughly 50% of Lys residues in the proteome are methylated. As a result, the true precision is expected to reside between the yellow and grey traces in Fig. 2A, which correspond to the estimated performance of MethylSight over the original (non-curated) test set and the high-confidence test set, respectively (see Methods for details). The performance of MethylSight was validated experimentally in Stage 4 using multiple reaction monitoring (MRM) mass spectrometry (Dhami et al., 2013; Liu et al., 2013) tailored for the predicted methylated Lys sites/peptides.

**Figure 2.**
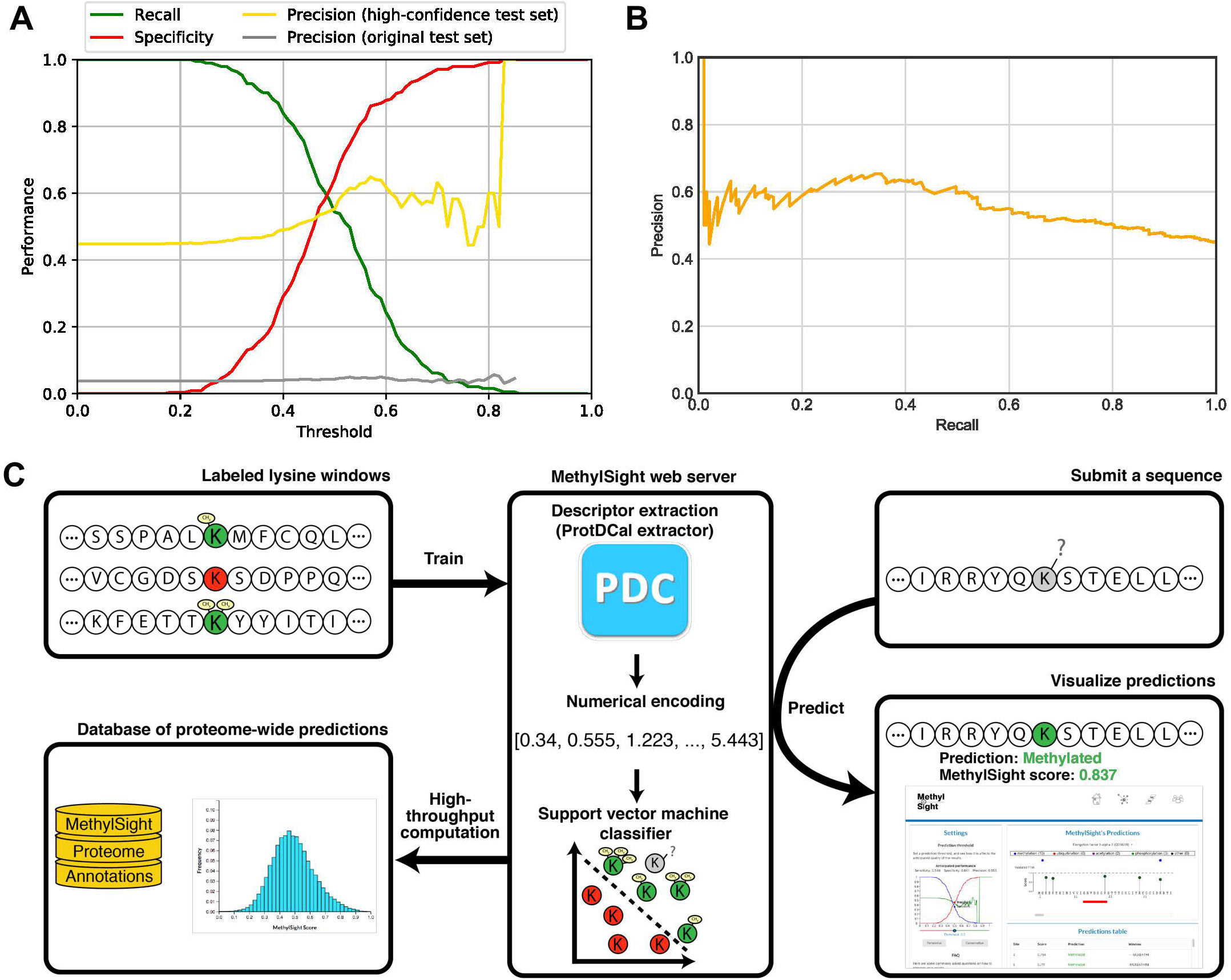
Performance of the Methylsight algorithm and use of the Methylsight software. (A) Performance measures of the MethylSight prediction of Lys methylation sites. The grey and yellow lines are pessimistic and optimistic estimators for true precision (see text for detailed description) such that the true precision falls between the two lines. (B) Precision-recall curve summarizing the anticipated precision of MethylSight as a function of the desired recall. (C) An overview of the MethylSight graphical user interface for methyl-Lys site prediction. See also Fig. S2 for a step-by-step guide in the use of the MethylSight web interface.

The model is subsequently evaluated in the hold-out test set, and the result contrasted with those from several available methylation prediction servers (Fig. S1). The performance of MethylSight is comparable to existing methods, with the exception of MethK (Lee et al., 2014) which achieved perfect precision on the test set at 60% recall (sensitivity). We note that MethK outputs a binary prediction (yes/no) as opposed to a confidence score. For this reason, there is a single precision-recall operating point with 60% recall, which inadvertently makes the method inflexible. In contrast, MethylSight outputs a confidence score (i.e., a measure of probability with a value between 0 and 1), and as such, enables the user to decide the recall level (Fig. 2C). Additionally, it cannot be guaranteed that the Kme sites used in our test set were not used to train MethK. Therefore, only a truly independent test set, using future data, will be able to fairly compare all methods. MethylSight is accessible via the webserver, http://methylsight.com, with the associated graphical user interface outlined in Fig. S2.

### MethylSight identifies novel Lys methylation sites on histones H1 and H2

Within the histone protein category, MethylSight predicted 70 Kme sites that have not been previously identified (Table S2). The majority of the new Kme sites is found on histone H1 with the remaining on H2A and H2B. Comparing to histones H3 and H4, in which nearly all Lys residues in their respective N-terminus have been found methylated, no Kme site has so far been identified in H2A and only a handful have been identified in H2B with low frequencies by MS-based high throughput (HTP) approaches (http://phosphosite.org). Similarly, the number of identified Kme sites in any given H1 variant is small compared to the large number of Lys residues contained in its sequence. Therefore, the prediction of the 70 new Kme sites suggests that histones H1 and H2 may be methylated on Lys residues as prevalently as H3 and H4.

Because it is relatively straightforward to isolate and purify histones, we decided to validate the predicted histone Kme sites by MS. To this end, we purified histones from the nuclear fraction of MCF7 cells (Fig. S3) (Schnitzler, 2001) and carried out MRM-MS analysis on the tryptic digested histone peptides using transitions specific to the predicted methylation sites (Table S3). Although our initial goal was to design MRM experiments for all 70 predicted Kme sites, this turned out to be impractical due to the high content of Lys/Arg residues on histones, which yields fragments too small for detection by MS upon digestion by trypsin (which cuts at the Lys/Arg residue). Nevertheless, using Skyline (MacLean et al., 2010), we were able to perform MRM-MS experiments for 50 predicted histone Kme sites and validated the existence of 45 sites in the MCF7 cells, some of which were found in multiple methylation states (i.e. mono-, di- or tri-methylation) (Table 2; Fig. S4). This corresponds to a validation rate of 90%, greatly surpassing the validation rate on the test set. This is remarkable given that all Kme sites are unlikely to be found in a given cell under a given condition. It is noteworthy that most of the validated Kme sites are on H1, a linker histone not known to harbor an abundance of methylation sites (Weiss et al., 2010). Furthermore, several novel Lys methylation sites were identified on the core histones H2A and H2B. The extremely high validation rate of the new Lys methylation sites on histones H1, H2A and H2B suggest that MethylSight is an effective algorithm by which to predict authentic Lys methylation events and that all histone proteins are subjected to extensive Lys methylation in cells.

**Table 2.**
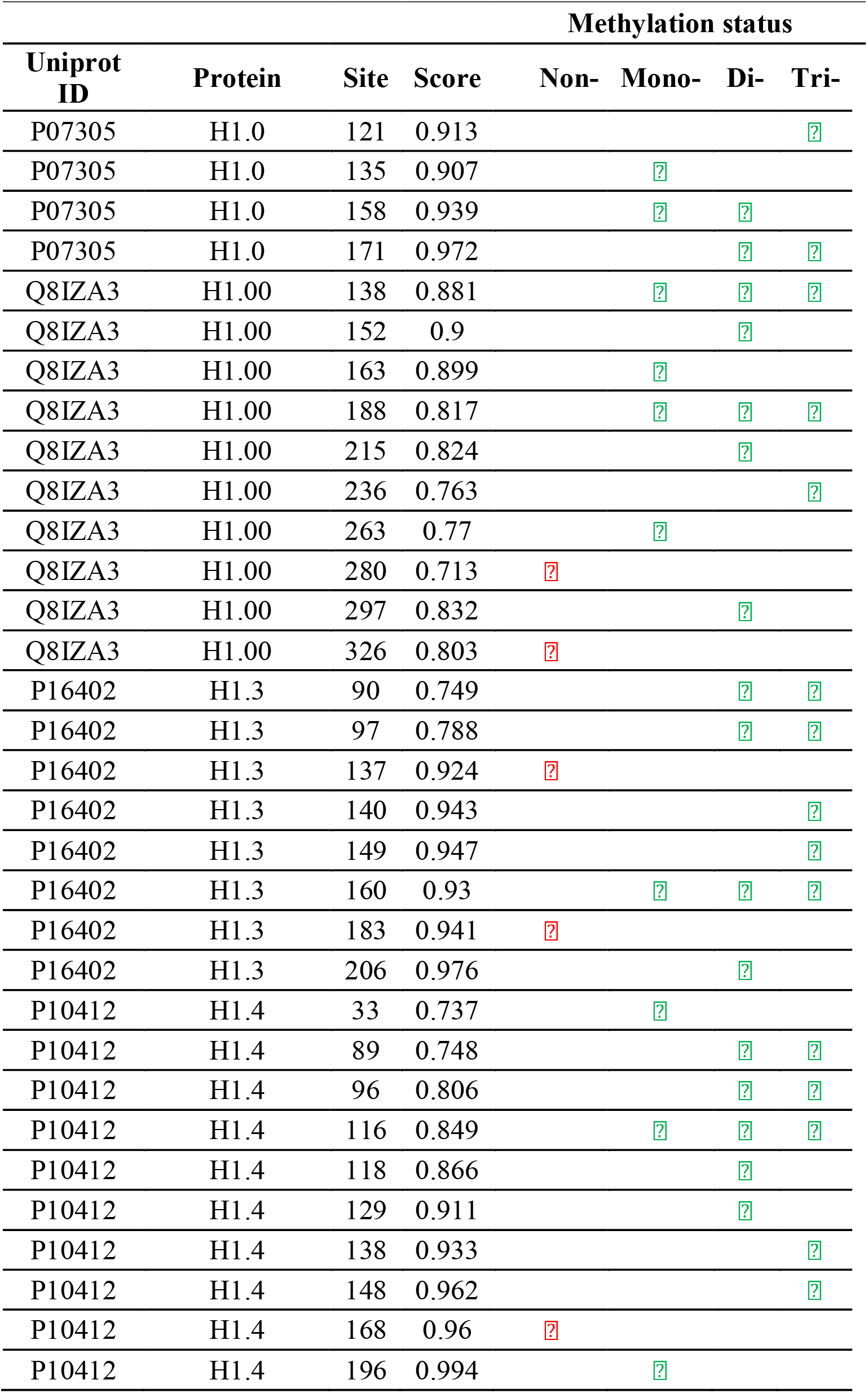
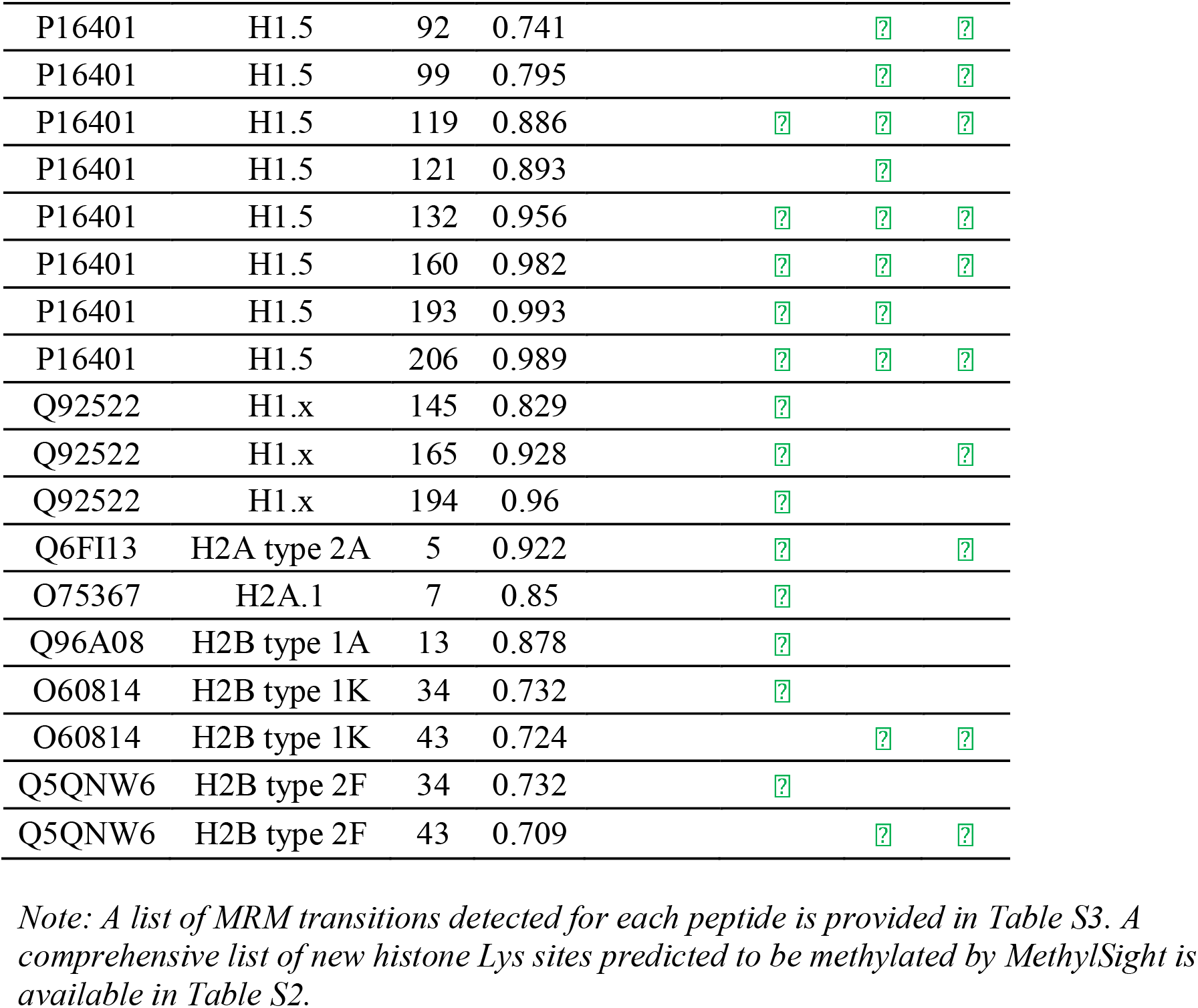
New histone Kme sites predicted by MethylSight and validated by MRM.

The MRM-validated new histone marks provided us with an independent test set to evaluate the performance of MethylSight against existing prediction algorithms such as MethK (Lee et al., 2014) and GPS (Deng et al., 2017). We found that all existing method, including MethK, showed poor recall to the 45 validated histone Kme sites (Table 3) even when the most liberal (lowest) decision threshold allowed by the corresponding software was used. Of note, MethK, which showed slightly better precision than MethylSight on the test set (Fig. 2), achieved a recall of 6.7% by correctly predicting only 3 of the 45 validated histone Kme sites. Therefore, MethylSight dramatically outperformed existing methods in accurately predicting new Kme sites.

**Table 3.**
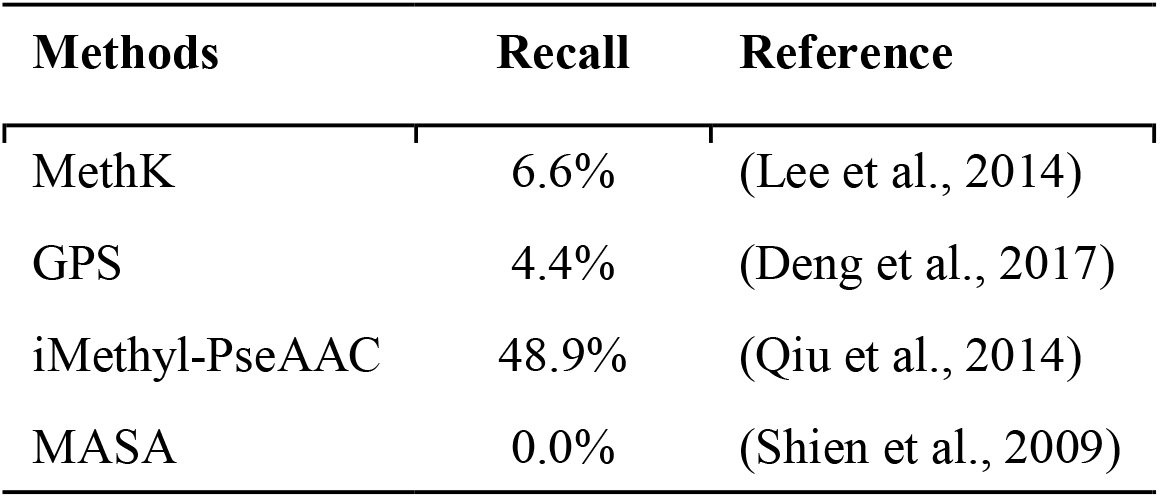
Performance of existing algorithms on the new histone Kme sites predicted by MethylSight and validated by MRM.

### H2BK43me2 is a novel substrate of KDM5b

Structural modeling shows that H2B methylation at Lys43 (i.e., H2BK43) is in close proximity to the phosphate backbone of the nucleosome DNA (Fig. S5), suggesting a potential role for this histone mark in nucleosome structure and/or function. We showed that H2BK43 could be di- or tri-methylated in MCF7 and mouse embryonic stem cells (mESCs) (Table 2; Fig. S6). To define the function of the H2BK43 methylation mark, we sought to first identify the corresponding Lys methyltransferase (KMT) or methyl-Lys demethylase (KMD). We therefore included the H2B peptides containing the K43me2 or K43me3 mark in an *in vitro* demethylation assay on synthetic histone peptides (Fig. S7). Using the MRM-MS strategy (Yocum and Chinnaiyan, 2009), we quantified the change in methylation for each histone peptide substrate in the presence of a recombinant Lys demethylase. We found that the H2BK43me2 peptide was the best substrate for KDM5b amongst the 15 histone peptides examined (Fig. 3A). Although KDM5b has been known to be a specific demethylase for the histone mark H3K4me3 (Xiang et al., 2007; Yamane et al., 2007), we found that the H2BK43me2 peptide was a superior substrate than either H3K4me2 or H3K4me3 for KDM5b, as ~80% of the former peptide was demethylated in 60 min compared to 20-30% for the latter (Fig. 3A). The remarkable activity of KDM5b towards the H2BK43me2 mark appeared to be specific for the dimethyl state because it showed no activity towards the trimethylated peptide. To characterize the kinetics of H2BK43me2 demethylation by KDM5b, we repeated the *in vitro* demethylation assay and took samples at different timepoints for MRM-MS analysis to monitor the progress of the enzymatic reaction (Fig. S8). The H3K4me3 and H2BK43me3 peptides were included as positive and negative controls, respectively. As shown in Fig. 3B, the H2BK43me2 peptide exhibited a fast kinetics of demethylation characterized with ~50% substrate turnover in 4 min, and by 30 min, the reaction was saturated (Fig. 3B). In contrast, only 8% of the H3K4me3 peptide was demethylated in 4 min, and 20% in 30 min. No significant change in the H2BK43me3 peptide was observed within the same time frame. These data indicate that the H2BK43me2 mark is a better substrate than H3K4me3 for KDM5b *in vitro*. To test whether the two peptides are competing substrates for KDM5b, we carried out the demethylation assay on the H2BK43me2 and H3K4me3 peptides mixed at an equal molar ratio. While both peptides were found demethylated, significantly less H2BK43me2 (<20%) than H3K4me3 (~40%) peptide remained after 60 min incubation with recombinant KDM5b (r.KDM5b) (Fig. S9). This indicates that KDM5b can use both the H2BK43me2 and H3K4me3 marks as substrates, yet it demethylates H2BK43me2 more efficiently than H3K4me3.

**Figure 3.**
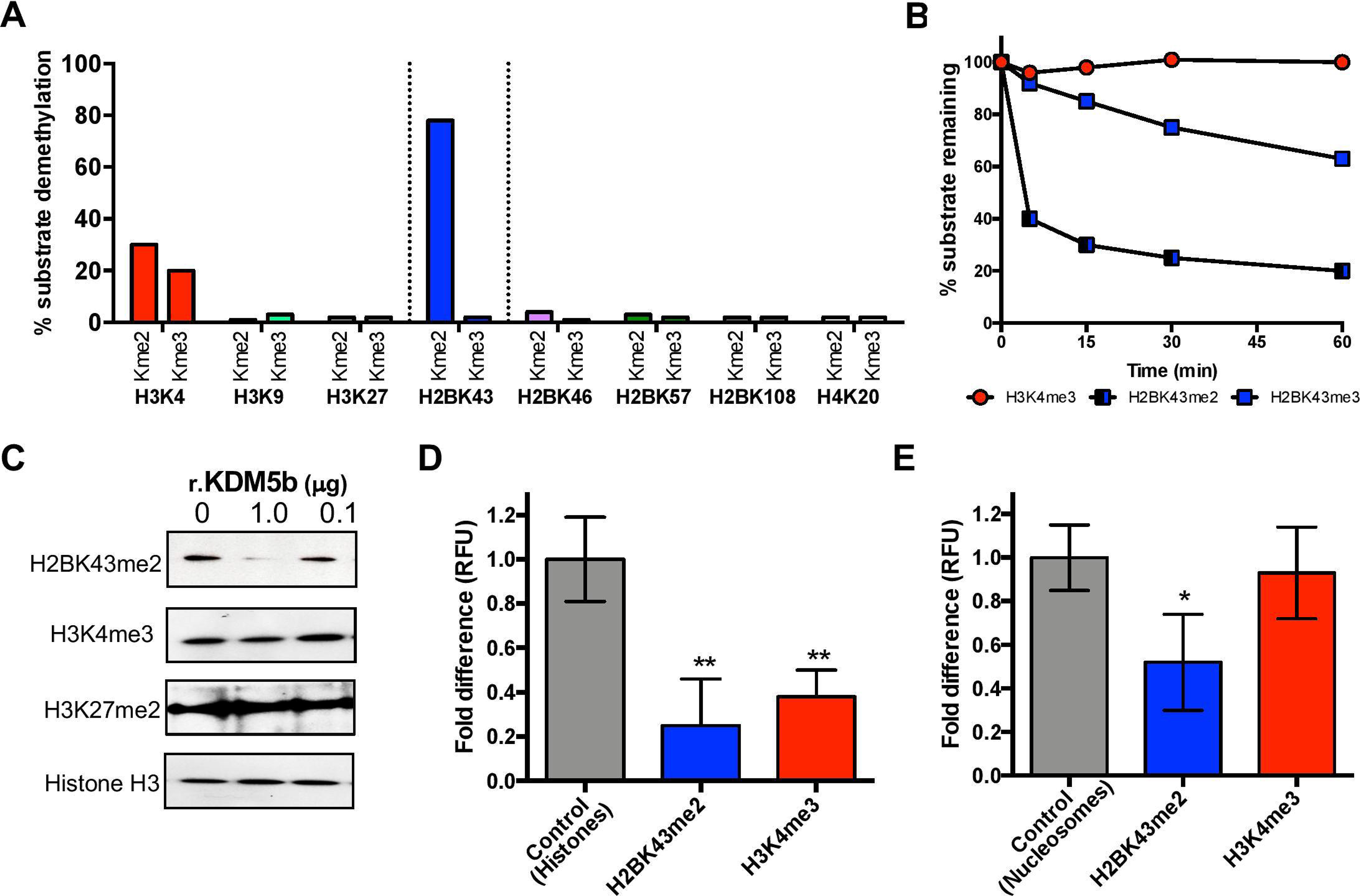
H2BK43me2 is a preferred substrate of KDM5b. (A) The activity profile of recombinant KDM5b (r.KDM5b) on histone peptides determined by in vitro demethylation assay. (B) H2BK43me2 is a preferred substrate for KDM5b in vitro. The amount of the remaining peptide substrate, H2BK43me2, H2BK43me3 or H3K4me3, was measured by MRM-MS at the 0, 5, 15, 30 and 60 minute of incubation with r.KDM5b and graphed as percentage of remaining substrate against time. (C) Recombinant KDM5b is capable of demethylating H2BK43me2, and, to a lesser extent, H3K4me3, but not H3K27me2 in the context of the nucleosome. Nucleosomes prepared from mESCs were subjected to demethylation reaction by r.KDM5b applied at the indicated amounts and the resulting products were separated on SDS-PAGE and immunoblotted for H3K43me2, H3K4me3, and H3K27me2, respectively. A pan-H3 Western blot was used to show the amount of total histone H3. (D) Immunoprecipitated (IP) KDM5b from mESC demethylates H2BK43me2 and H3K4me3 in purified nucleosomes. The efficiency of demethylation by KDM5b of H2BK43me2 or H3K4me3 was measured by a microplate-based fluorescence assay. Shown on the graph is the amount of either histone mark in the presence of KDM5b relative to that in the absence of the enzyme, set at 1. RFU, relative fluorescence unit. (E) Both the H2BK43me2 and H3K4me3 marks on bulk histones could be demethylated by endogenous KDM5b IP’ed from mESCs *, p<0.05; **, p<0.001, Student’s t-test (n=6).

### KDM5b specifically demethylates H2BK43me2 *in vivo*

To determine whether H2BK43me2 is a substrate of KDM5b *in vivo*, we repeated the demethylase assay on nucleosomes and histones. Given the role of KDM5b in stem cell differentiation (Dey et al., 2008; Xhabija and Kidder, 2019), we purified nucleosomes from the mESCs and incubated them with r.KDM5b. The demethylation of nucleosomal H2BK43me2 was assessed by Western blotting. In parallel experiments, we analyzed the demethylation of the H3K4me3 and H3K27me2 marks (as a negative control). An anti-histone H3 Western blot was included as a loading control. The anti-H2BK43me2 antibody used in this study recognized primarily the H2BK43me2 mark, and, to a much lesser extent, H2BK43me1, but showed no binding to other histone marks on a peptide spot array (Fig. S10). In comparison, the anti-H3K4me3 antibody bound primarily to the H3K4me3 mark and to other histone marks in a marked reduced degree (Fig. S10). Consistent with results from the MRM-MS analysis, incubation of the nucleosome with 1 µg r.KDM5b resulted in a marked reduction in the H2BK43me2 level, but only a slight decrease in H3K4me3 and no change in the H3K27me2 level (Fig. 3C). These data indicate that r.KDM5b preferentially demethylates the H2BK43me2 mark on the nucleosome.

Mammalian proteins often contain post-translational modifications that are lost when expressed in bacteria. To confirm that the observed activity of the recombinant KDM5b on the H2K43me2 mark recapitulates the specificity of the enzyme in mammalian cells, we immunoprecipitated endogenous KDM5b from the mESCs and employed it in demethylation assays using either purified histones or nucleosomes as substrates. The endogenous KDM5b exhibited activity towards H2BK43me2 present on histone H2B (Fig. 3D) and was able to demethylate H2BK43me2 significantly more efficiently than H3K4me3 in the context of the nucleosome (Fig. 3E). These data demonstrate that the H2BK43me2 mark is a better substrate than H3K4me3 for KDM5b *in vivo*.

### The global level of H2BK43me2 undergoes dynamic changes during mESC differentiation

To the best of our knowledge, the function of mammalian H2BK43me2 has not been characterized to date. Because KDM5b plays an important role in stem cell differentiation (Dey et al., 2008; Schmitz et al., 2011), it is likely that the H2BK43me2 mark also plays a role in this process. To explore this possibility, we first examined whether KDM5b is regulated during the course of neuronal differentiation of mESCs. The KDM5b protein level declined on day 5 of differentiation when the cells were committed to the neuronal lineage; and by day 7, it dropped to a level below detection by Western blot (Fig. 4A). The dynamic expression of KDM5b during neuronal differentiation of mESCs suggests that the abundance of its substrate would follow an opposite trend. To examine whether this was the case with H3K4me3 and H2BK43me2, we immunoblotted for these marks in mESCs collected at days 0, 5 and 7 of neuronal differentiation. While the level of H3K4me3 remained unchanged based on the Western blot, a marked increase in the global level of H2BK43me2 was observed on days 5 and 7 (Fig. 4B).

**Figure 4.**
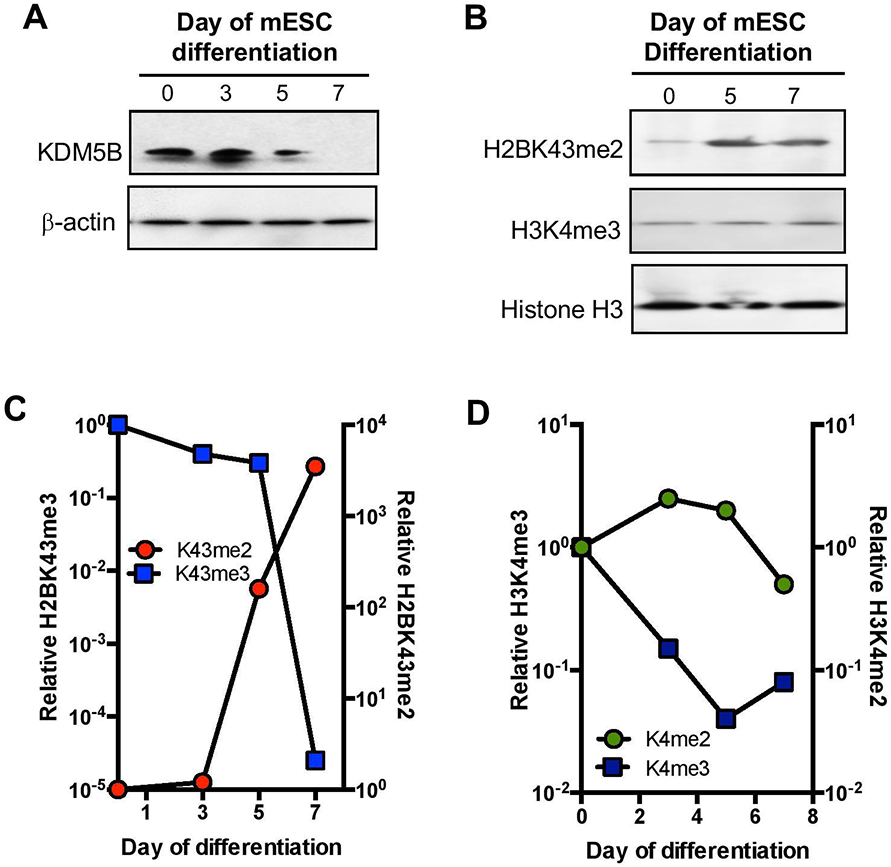
H2BK43me2 undergoes dynamic changes during mESC neuronal differentiation. (A) The protein level of KDM5b decreases in mESCs during neuronal differentiation. (B) A Western blot showing that the global level of H3K4me3 remains constant whereas that of H3K4me2 increases markedly during mESC neuronal differentiation. (C) H2BK43me2 is a highly dynamic histone mark. The global H2BK43me2 level increases by several orders of magnitude at the late timepoints (i.e. day 7) of neuronal differentiation. A reversed kinetic pattern was observed for H2BK43me3. (D) Changes in global H3K4me2 and H3K4me3 levels during neuronal differentiation of mESCs measured by MRM-MS. Note the difference in scale between panels C and D.

To quantify the dynamic changes of the two KDM5b substrates during ESC differentiation, we used MRM-MS to determine the abundance of H3K4me2 and H3K4me3 and H2BK43me2 and H2BK43me3 in cell number-matched mESC samples taken on days 0, 3, 5, and 7 of neuronal differentiation (Tables S4 and S5). The H2BK43me2 and H2BK43me3 marks exhibited marked dynamic changes during mESC differentiation. Specifically, the level of H2BK43me3 decreased gradually in the early stages of differentiation, but it dropped sharply between days 5 and 7 (Fig. 4C). By day 7, an approximately 10,000-fold decrease in the global level of H2BK43me3 was observed, relative to the level on day 0. Conversely, the H2BK43me2 mark underwent a dramatic increase between days 3 and 7, and by day 7, it reached a level approximately 10,000-fold greater than that of day 0 (Fig. 4C). In contrast, the levels of the H3K4me2 and H3K4me3 marks underwent much smaller changes during the same time frame (Fig. 4D). Collectively, these data demonstrate a reverse correlation between the global levels of KDM5b and H2BK43me2 during mESC neuronal differentiation and identify H2BK43me2 as a highly dynamic histone mark in this process.

### Prediction of the human methyl-Lys proteome

To estimate the breadth of Lys methylation in the human proteome, we used MethylSight to predict proteins harboring high-confidence Lys methylation sites. A confidence score of 0.7 was chosen for the prediction which corresponds to 95% specificity (Fig. 2A). At this confidence score, 48,459 Lys residues contained in 11,259 proteins were predicted to be methylated (Table S6). The predicted human methyl-Lys proteome comprises 11,206 non-histone proteins and 53 histone proteins of which 2,008 have been annotated in PhosphositePlus, 9,251 are novel.

To gain insights into the functions of Lys methylation, we carried out a GO analysis (Ashburner et al., 2000) on both the known (as annotated at PhosphoSitePlus) and the MethylSight predicted methyl-Lys proteome. The GO analysis identified 19 biological processes or molecular functions that are enriched in the predicted but are not in the known methyl-Lys proteome (Fig. 5A). Of note, statistically significant enrichment (p<0.05) of Lys methylated proteins was observed in biological processes that include ribosome biogenesis, protein translation, RNA processing, DNA metabolism, chromatin organization, cell cycle process, and cellular complex and organelle organization (Fig. 5A).

**Figure 5.**
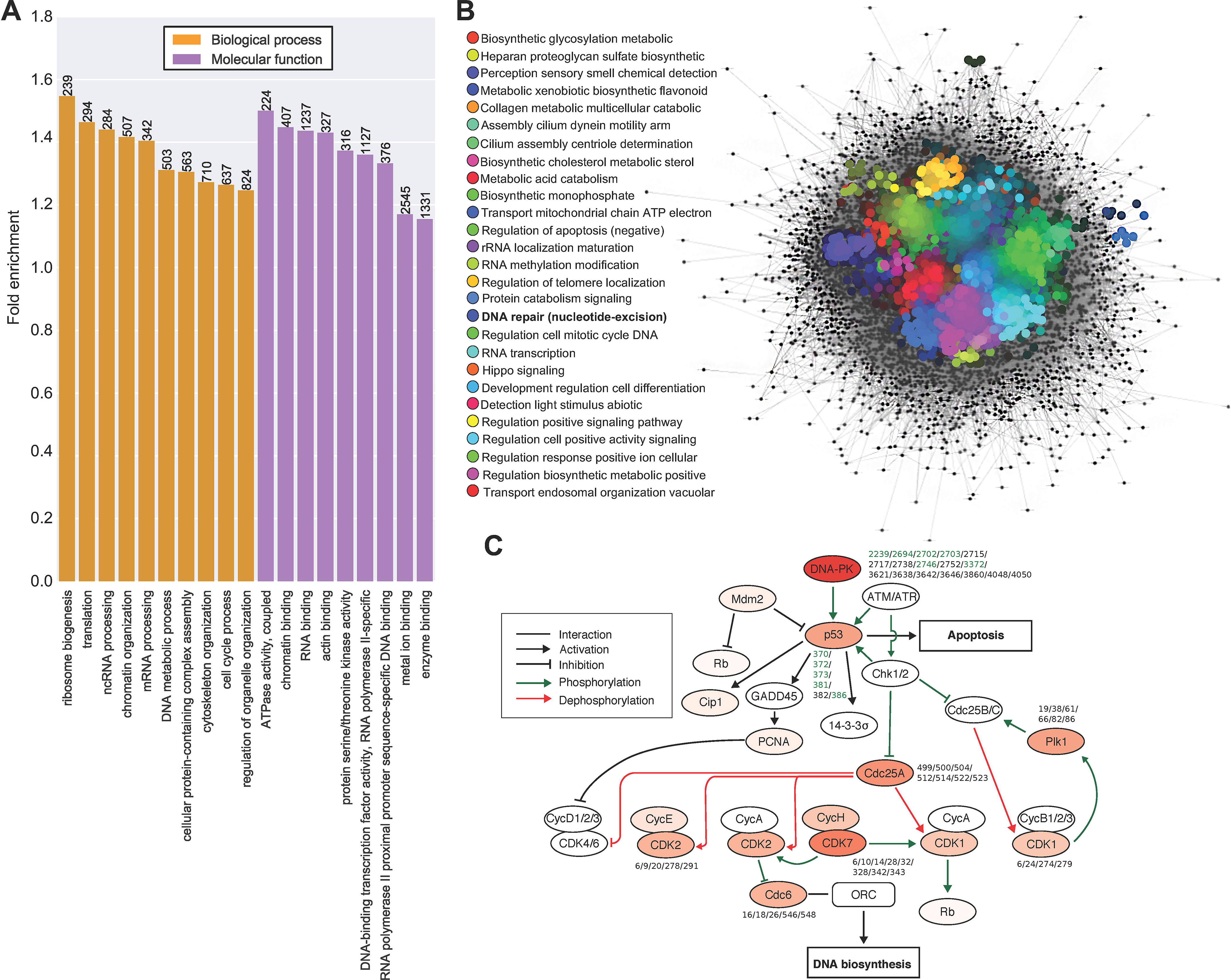
The predicted human methyl-Lys proteome. (A) Gene Ontology (GO) annotation of the biological processes and molecular functions enriched in the MethylSight predicted methyl-Lys proteome. The number of methylated proteins associated with a given GO term is indicated. (B) A network map of both known and MethylSight predicted Lys-methylated proteins. The map were created by STRING with a confidence cut-off of 0.4. SAFE annotation of the network with 27 GO biological process terms shows the interaction landscape of the predicted methyl-Lys proteome. (C) A subset of the cell cycle KEGG pathway (KEGG ID: hsa04110) annotated with MethylSight at a 0.7 confidence threshold. Proteins are colored on a yellow-to-red scale, with darker shades indicating a higher degree of methylation. Validated Lys methylation sites are identified in green, while predicted methylation sites are shown in black.

To better understand how the predicted methyl-Lys proteome is organized, we carried out a spatial analysis of functional enrichment (SAFE) (Baryshnikova, 2016), a method developed for the purpose of annotating biological networks and examining functional organization. The SAFE analysis was carried out on a network comprising both known and MethylSight-predicted Kme-modified proteins. The methyl-protein network created using STRING (Szklarczyk et al., 2019) with a confidence cut-off of 0.4, consisted of 11,311 nodes and 429,160 edges and were organized in 2D space in Cytoscape (v.3.7.2) (Shannon et al., 2003). SAFE annotation of the network with 27 GO biological process terms identified the core of the interaction landscape of the predicted methyl-Lys proteome that is densely connected. Of note, significant enrichment of methyl-Lys was observed in a variety of different cellular processes including RNA transcription, metabolism, cell cycle regulation, DNA damage response and signal transduction (Fig. 5B).

To provide a closer view of the methyl-Lys proteome, we examined the predicted methyl-Lys protein network in the context of DNA damage response (DDR) and cell cycle progression. Although previous studies have identified multiple Kme sites in p53 and DNA-PK (West and Gozani, 2011; Liu et al., 2013) that play important roles in regulating DDR and apoptosis, MethylSight predicted numerous novel Kme sites that likely exist in DNA-PK and protein kinases (such as cyclin-dependent kinases or CDKs) that are involved in cell cycle regulation (KEGG ID: hsa04110). Of particular interest is the predicted high-degree of methylation occurring on DNA-PK, Cdc25A, CDK7, CDK2 and CDK1. It is likely that Lys methylation plays a role, not only in regulating DDR and cell cycle progression, but also in the crosstalk between these cellular processes (Fig. 5C).

## DISCUSSION

Though much effort has been devoted to the investigation of protein methylation, systematic identification of Lys methylation by experimental approaches has remained a great challenge. To overcome this bottleneck, a number of *in silico* methods have been developed for the prediction of Lys methylation (Chen et al., 2006; Deng et al., 2017; Hu et al., 2011; Lee et al., 2014; Qiu et al., 2014; Shao et al., 2009; Shi et al., 2012a; Shien et al., 2009). However, these methods are limited by the small amount of experimentally validated Kme sites available at the time for training. Moreover, their power in predicting new Kme events has not been rigorously tested. We developed MethylSight, based on the largest number of annotated Kme sites available for training, to speed up the pace of discovery for potential Lys methylation sites with increased prediction accuracy. Indeed, the MethylSight algorithm, an SVM-based prediction tool, was trained using alignment-free features to encode the sequence-based information for the thousands of Kme sites that have been experimentally annotated.

At first glance, the overall performance of MethylSight appears to be comparable to competing methyl-Lys predictors (Fig. S1). However, our test set was prepared without prior knowledge of the specific sites used in the training set of competing methods. It is therefore highly probable that a large number of Kme sites used to benchmark the competing predictors were, in fact, used to train them. This heavily biases the comparison in favour of competing predictors. Indeed, we have shown here that MethylSight is powerful in discovering novel methyl-Lys modification events with remarkable accuracy. Although histone Lys methylation has been extensively investigated in the past two decades, MethylSight predicted the existence of 70 new histone Kme sites, suggesting that much remains to be learnt about histone Lys methylation. That 90% of the predicted Kme sites in the histones H1 and H2A/B were subsequently validated in cells, suggests that MethylSight is a highly effective algorithm for the identification of authentic Kme sites. Using these experimentally validated Kme sites, we re-evaluated the performance of existing prediction methods and found the latter performed poorly (Table 3). Our observation that the previous methods exhibited good performance over our test set, but showed poor recalls over the validated new sites, suggests that they may be overfit to the training data and thus of limited use as general approaches to predict the methyl-Lys proteome.

Our work also establishes a robust pipeline for systematic characterization of Lys methylation by combining MethylSight prediction with experimental validation using targeted mass spectrometry approaches such as MRM/SRM (Selected Reaction Monitoring) (Dhami et al., 2013; Liu et al., 2013). While the functions of the new histone Kme sites remain to be defined, it is tempting to speculate that H1 methylation on multiple Kme sites simultaneously may facilitate its function as the linker histone by stabilizing it and preventing its association with DNA. Furthermore, our data suggests that while H3K4me3 is a general histone mark for transcription activation (Liang et al., 2004), H2BK43me2 may be a mark specific for KDM5b targeted genes (Klein et al., 2014). These two marks may work together to provide an important epigenetic control of the transcriptional program during ESC differentiation. It would be interesting to determine in subsequent studies whether the H2BK43me2 mark regulates gene expression on a genome-wide scale in the context of stem cell differentiation or other cellular processes.

Besides histones, MethylSight predicted the methyl-Lys proteome to consist of ~50,000 methyl-Lys sites; an approximate 10-fold increase over the <5,000 sites currently annotated in the PhosphoSitePlus database. The size of the predicted methyl-Lys proteome is comparable to that of the annotated phospho-Tyr proteome. This is not surprising given the comparable number of Lys methyltransferases (KMTs) and Tyr kinases (Biggar and Li, 2015; Corwin et al., 2017; Paul and Mukhopadhyay, 2004). However, the functions of the methyl-Lys proteome and phospho-Tyr proteome are markedly different. While a large proportion of Lys methylated proteins are found in the nucleus, the majority of Tyr phosphorylated proteins are cytosolic. Nevertheless, cross-talks between these two prevalent post-translational modifications are bound to happen. For example, the fortuitous discovery of MAP3K2 methylation at K260 by SMYD3 was shown to be instrumental in the activation of oncogenic Ras/Raf/MEK/ERK signalling and the progression of Ras-driven lung cancers (Mazur et al., 2014). This example highlights the importance of developing tools that are able to accurately identify new Lys methylation sites for their functional annotation.

To gain functional insight into the predicted methyl-Lys proteome, we performed a GO annotation analysis to identify in Lys methylation significantly enriched in cellular processes. The highest enrichment was found for GO terms related to ribosome biogenesis and translation. These results align well with those from recent studies showing that Lys methylation of elongation factors is a prevalent PTM. In fact, it has been shown that methylation of the yeast elongation factor 2 on K509, which is conserved in humans, modulates its affinity for the 40S subunit of the 80S ribosome (Zhang et al., 2014). In a more recent study, Lys methylation of human elongation factor 1-alpha 1 was shown to mediate such processes as cytoskeleton organization, rRNA aminoacylation and ribosome biogenesis in a methylation site-dependent manner (Hamey et al., 2017).

Given that our GO term enrichment analysis showed significant enrichment for Lys methylation in proteins associated with the cell cycle and DNA damage repair processes, we elected to identify potentially methylated proteins in the cell cycle KEGG pathway. Of note, DNA-PKcs, a member of this pathway, was predicted to be methylated on 11 new Lys sites. At the time of writing, the PhosphoSitePlus database tabulated 6 methyl-Lys sites within DNA-PKcs, of which the methylation on K1150, K2746 and K3248 has been shown to play a role in its interaction with the chromodomain of HP1β (Liu et al., 2013). Five of the potential methylation sites identified by MethylSight in DNA-PKcs are located in its kinase interacting protein (KIP)-binding region. It is tempting to speculate that Lys methylation within this region would modulate the interaction of DNA-PK with HP1β. This interaction is believed to be important for telomere end-capping as it is necessary to recruit DNA-PKcs to the telomere (Khadka et al., 2014). Given that DNA-PKcs has emerged as a potential therapeutic target in cancer (Mohiuddin and Kang, 2019), there is significant incentives to validate these potential methylation sites and identify their associated KMT/KDM in subsequent studies.

The MethylSight algorithm (accessible via http://methylsight.com) will provide a robust tool and valuable resource for the discovery and characterization of novel Lys methylations occurring on both histone and non-histone proteins. When combined with high throughput experimental approaches such as mass spectrometry, MethylSight would enable systematic annotation of Lys methylation and characterization of the dynamic changes in protein methylation associated with cellular functions. Furthermore, the discovery of, and mechanistic insights into, new Lys methylation events will undoubtedly pave the way for the future development and therapeutic application of “epi-drugs” in the treatment of cancer and a host of other diseases (Arrowsmith et al., 2012).

## ACKNOWLEDGEMENTS

This work was supported by a National Science and Engineering Research Council (NSERC) Canada postgraduate scholarship to FC, NSERC Canada Discovery grants (to KKB, JRG and SSCL, respectively) and by a Canadian Institutes of Health Research (CIHR) grant (to SSCL). SSCL holds a Canada Research Chair (Tier I) in Molecular and Epigenetic Basis of Cancer.

## AUTHOR CONTRIBUTIONS

K.K.B., H.L., L.S., A.C., J.C., H.A., K.F., M.G., and Q.F. conducted the experiments, F.C. and Y.B.R.B. developed the MethylSight algorithm, K.K.B., C.W., W.S., J.R.G., and S.S.C.L. designed the experiments, and K.K.B., F.C., J.R.G., and S.S.C.L. wrote the paper.

## METHODS AND MATERIALS

### LEAD CONTACT AND MATERIALS AVAILABILITY

Further information and requests for resources and reagents should be directed to and will be fulfilled by the Lead Contact Shawn Li.

#### Detailed Method for MethylSight

##### Dataset preparation

All Lys methylation data were retrieved from the PhosphoSitePlus curated methylation dataset (Hornbeck et al., 2015). This dataset comprises positive methylation sites only. As such, Lys occurring within methylated proteins, but not previously reported to be methylated, were assumed to be non-methylated. All Lys residues in the dataset were processed into 71 amino acid windows centered around the Lys residue (Shi et al., 2015b), padding with Ala residues wherever the Lys is too close to the N- or C-terminus of the protein.

To reduce the overall redundancy of the dataset, the windows were clustered at a 90% identity level with CD-HIT, and a representative of every cluster was sampled (Fu et al., 2012). Where a positive and a negative site are found sharing an identity above this threshold, the negative one was removed. Subsequently, a random sample corresponding to 10% of the filtered dataset was set aside to build a hold-out test-set to assess the generalizability of our model over unseen data. The remaining 90% of the data was further filtered at a 70% identity level, before randomly sampling 80% and 20% of the data to build training and validation sets, respectively.

To mitigate the fact that many “assumed-to-be” negative sites may be undiscovered or mislabeled positives, we sought to further refine our test-set in order to derive more accurate insight into the performance of our classifier. To this end, we opted to remove sites from our initial test set that are either known to be acetylated, ubiquitinated or sumoylated, since these sites are demonstrably available to enzymes and there is a known correlation between sites that are modified in multiple ways. In addition, we used NetSurfP 1.0 (Petersen et al., 2009) to identify solvent accessible Lys residues. We considered all Lys residues with a relative solvent accessibility lower than 0.2 to be buried, and only retained these sites, given that they are unlikely to be accessible to a KMT. Collectively, this would minimize the chance that any of the remaining sites are false negatives. Table 2 lists all the datasets used in this study.FlyF

##### Feature extraction

We used the ProtDCal software (Ruiz-Blanco et al., 2015) to generate a feature vector for each Lys-centered window in the dataset. The feature vector consists of a set of scalar values computed from tabulated physico-chemical characteristics of the amino acids using a variety of aggregation operations to capture the properties of amino acids and their neighborhood. A total of 3720 descriptors can be derived from a potential methylation site, although it is possible to limit this to a subset by means of an IDL file specifying which descriptors should be computed, once feature selection is performed. In this case, 12 amino acid properties were used to numerically encode the physical-chemical characteristics of a methyl-Lys residue. These properties are found in the AAindex database (Kawashima and Kanehisa, 2000) and are also described in the ProtDCal documentation. Fourteen residue groups are used based on either side chain structure or using specific residue positions within the input sequence window. Two modification operators for capturing vicinity information and twelve aggregation operators ultimately transform the property vector of each amino acid group into the final scalar features. The project files with the lists of indices, groups, modification and aggregation operators as well as other parameters for the calculations are provided at https://github.com/flexplicateur/MethylSight.

##### Feature selection

Feature selection began with information gain (IG) analysis, which retains only those features whose distribution across all sites in the training data correlates with class label. All the attributes with a non-zero IG value were extracted in this step. Subsequently, an unsupervised redundancy filter was applied, using a single-linkage clustering algorithm with the Spearman correlation coefficient as the similarity measure. Features exhibiting pair-wise correlation above 0.9 were clustered together and only one representative feature from each cluster was kept. Ultimately, the supervised WrapperSubsetEval method, implemented in Weka 3.7.11 (Hall et al., 2009), was used to extract an optimum subset of features for modelling. This method was configured using a Genetic Search for exploring the feature space and potential feature sets are evaluated using the classification F-measure of the positive class in 5-fold cross-validation tests using support vector machine (SVM) classifiers with a linear kernel. The cost-sensitive sequential minimal optimization (SMO) algorithm (Cai and Cherkassky, 2012) was used to train all SVM classifiers in this work. The cost matrix reflected the relative class imbalance in the data, such that the false negative error cost is equal to the number of negative instances and the cost of false positive errors was fixed at the number of positive instances in the data (Table 1).

##### Support vector machine classifier

Support vector machines (SVMs) are widely used machine learning models for binary classification. They are widely used in bioinformatics for a variety of tasks including protein-protein interaction prediction (Guo et al., 2008; Romero-Molina et al., 2019), protein function prediction (Cai et al., 2003), sub-cellular localization prediction (Chou and Cai, 2002), and post-translation modification prediction (Shi et al., 2015b). Training instances are projected in a *k*-dimensional Euclidean space according to the *k* descriptors, and the SVM model parameters are optimized to find the hyperplane, or boundary, that best separates positive instances from negative instances, i.e. with the largest margin. In addition to these parameters learned from the training data, SVMs rely on hyperparameters specified by the user and that are not directly inferred from the training data. These hyperparameters control the penalty associated with a misclassification, among others.

In this study, we use SVM models to select the most relevant descriptors for a given Lys-centered window and predict how likely the Lys is to be methylated. The cost-sensitive sequential minimal optimization (SMO) algorithm (Cai and Cherkassky, 2012) was used to train all SVM classifiers in this work. We used a cost matrix reflective of the relative class imbalance in the data in order to prevent bias towards the negative class, which is overwhelmingly more prevalent.

##### Training and validating the support vector machine predictor

Following a series of supervised and unsupervised steps taken to determine the optimal feature set (described in the Supplemental Materials), we performed a grid search to optimize the SVM model hyperparameters. The hyperparameters were varied iteratively, and the set of hyperparameters producing the best prediction accuracy in 5-fold cross-validation over the training set, as well as on the validation set (see Table 1) were selected. The F-measure, precision, and recall were used to assess performance. The anticipated performance of the predictor upon deployment was assessed using the hold-out test set of randomly sampled sites previously described consisting of 193 validated positive sites, and 237 “high-confidence” negative sites.

##### Functional annotation of the predicted methyl-Lys proteome

To functionally annotate the biological functions enriched in the dataset of known and predicted human Lys methylation sites, we initially used Gene Ontology (GO) enrichments to identify biological processes enriched in Lys-methylated proteins. The gene over-representation analysis was carried out in PANTHER (Mi et al., 2019), using Fisher’s exact test and the Bonferroni correction for multiple testing with a significant level of p<0.05. Complementary to this analysis, spatial analysis of functional enrichment (SAFE), a method for annotating biological networks and examining functional organization, was carried out on our dataset of known and predicted human Lys methylation sites using default settings and visualized using Cytoscape v.3.7.2 (Baryshnikova, 2016).

#### Experimental Methods Details

##### Antibodies

Mouse anti-H2BK43me2 was created via service by YenZym antibodies (San Francisco CA, USA). The specificity of the anti-H2BK43me2 antibody to detect distinct methylation states (i.e., Kme1, 2, or 3) were confirmed by immunoblotting against dot blots that contained synthesized H2B peptides that corresponded to a central K43 methylation site (YSVYVY**Kme**VLKQVH) either without the methyl-modification or in the presence K43me1, 2 or 3 methylation (Fig. S10). Rabbit anti-H2B (cat# 07-371) and anti-H3 (cat# 07-690) antibodies were purchased from EMD Millipore. Other methylation-specific histone antibodies used in this study include rabbit anti-H3K4me3 (EMD Millipore, Cat# 04-745), and rabbit anti-H3K27me2 (at 1:1000 dilution; Abcam, cat# ab24684).

##### Cell culture and neuronal differentiation

MCF7 cells were culture in DMEM (Sigma Aldrich) supplemented with 10% FBS (Wisent), 2mM L-glutamine (Gibco), 100U/ml penicillin-streptomycin (Gibco), 2mM sodium pyruvate (Gibco) at 37 °C humidified incubator with 5% CO_2_. Mouse embryonic stem cells E14 were cultured using feeder-independent protocol (Bibel *et al*., 2007). In brief, mESCs were cultured on 0.1% gelatin-coated tissue culture treated dishes in DMEM (Sigma Aldrich) supplemented with 15% embryonic stem cell grade FBS (Wisent), 2 mM sodium pyruvate (Gibco), 2 mM L-glutamine, 100 U/ml penicillin-streptomycin (Gibco), 0.1 mM 2-mercaptoethanol (Sigma Aldrich), 1X non-essential amino acid (NEAA) supplement (Gibco) and 1000 U/ml LIF (Stem cell biotechnology), at 37 °C humidified incubator with 5% CO_2._ Cells passaged at sub-confluent stage of 70% to avoid stem cell colonies touching each other. To passage, cells were first washed with PBS with 0.1mM EDTA twice. Typically, 1 ml of 0.1% Trypsin was added to a 10 cm dish with 5 min incubation at 37°C. Complete dissociation of clumps should be achieved to avoid spontaneous differentiation after passage. For neuronal differentiation, mESCs were grown in mESC media until cells became highly proliferative (i.e., doubling time less than 20 hr). Highly proliferative mESCs were lifted and 4 × 10^6^ cells were seeded in bacterial Petri dish with 10 ml embryoid body media containing DMEM, 1% FBS (Gibco), 1X B-27 supplement (Gibco), 2mM L-glutamine, 1X NEAA supplement, 100 U/ml penicillin-streptomycin and 2 mM sodium pyruvate. Cells were left untouched for 48 hr to allow formation of embryoid bodies. Embryoid bodies were then plated onto 1% gelatin coated tissue culture treated dishes (1:4 ratio) with neuronal differentiation media consisting of 5% FBS, 1X B27 supplement, 2 mM L-glutamine, 100 U/ml penicillin-streptomycin and 2 mM sodium pyruvate. Cells were maintained in this condition for further differentiation days.

##### Mass Spectrometry

To validate the status of predicted methylation sites, isolated histone proteins were digested with trypsin and the digest was analyzed by positive ESI LC-MS/MS on a triple quadrupole mass spectrometer (4000 QTRAP, Applied Biosystems Inc.) utilizing Q3 as a linear ion trap. A nanoAcquity UPLC system (Waters) equipped with a C18 analytical column (1.7 µm, BEH130, 75 µm×250 mm) was used to separate the peptides at the flow rate of 300 nl/min and operating pressure of 8000 psi. Peptides were eluted using a 62 min gradient from 95% solvent A (H_2_O, 0.1% formic acid) and 5% B (acetonitrile, 0.1% formic acid) to 50% B in 41 min, 6 min at 90% B, and back to 5% for 10 min. Eluted peptides were directly electrosprayed (Nanosource, ESI voltage +2000V) into the MS. The instrument was set to monitor up to 200 transitions in each sample with a dwelling time of at least 25 msec/transition.

For multiple reaction monitoring MS (MRM-MS), the *in silico* protease digest patterns (i.e. to generate precursor ions) and the corresponding MRM transitions were compiled using the Skyline™ software (MacLean et al., 2010). Transitions that are larger than the precursor ion was selected based on the Skyline predictions and the specific b/y ions that allow unambiguous identification of the methylated Lys site were included and used in the isolation list. Positive identification of a new methylation site required the successful detection of at least three transition ions. All transitions used to identify methylation sites are listed in Table S3.

##### Histone extraction

We followed the histone extraction strategy from Shechter *et al* with minor modifications (Shechter et al., 2007). Cells were first collected by washed by ice-chilled PBS twice to remove FBS completely. Typically, 1 × 10^7^ cells were lysed with 5 ml ice-chilled lysis buffer containing 0.5% NP40 (Sigma Aldrich), 10 mM Tris-HCl [pH 7.6], 140mM NaCl, 1.5 mM MgCl_2_, 1 mM PMSF, on ice for 10 min with occasion inversions. Nuclei were pelleted by centrifugation at 2100 rpm, 4°C for 5 min. A total of 1 ml of 0.4N H_2_SO_4_ was added to the pellet and then incubated for 1 hr on ice. Supernatant was collected by centrifugation at 10000 rpm, 4°C for 5 min. In a new Eppendorf tube, mix 250 μl of TCA with the supernatant for about 30 min to precipitate the histone proteins. Precipitates were pelleted by centrifugation at 14000 rpm, 4°C for 10 min. Pelleted histones were firstly washed by acidified acetone (−20°C) then twice with 100% acetone (−20°C). Lastly, the pellet was air dried and resuspended in water. Quality of histone was assessed by SDS-PAGE and Coomassie staining.

##### Histone demethylase assay (HDM)

All histone demethylase reactions, including peptide, histone and nucleosome substrate assays, were performed as described in Lee et al (Lee et al., 2007) using demethylation buffer (50 mM HEPES [pH 8.0], 100 mM NH_4_SO_4_, 1 mM alpha-ketoglutarate, 2 mM ascorbate, 5% glycerol, and 0.2 mM PMSF). The peptide HDM reaction was prepared on ice and a time zero sample was removed. The samples were then incubated with r.KDM5b at 37°C and samples were removed at 5, 30, 60, 90, 120 and 360 min. For the synthetic H2BK43 peptide (Biotin-AHX-YVY**Kme2**VLKQVHPDT) 15 transitions were monitored (M→ y_10_-y_4_). For H3K4 synthetic peptide (Biotin-AHX-ART**Kme3**QTARKS) 20 transitions were monitored (M→ y_8_-y_2_). The peak area, being proportional to the quantity of a given peptide, was used for its relative quantification.

##### Quantitative HDM assay

The buffer conditions and procedures were essentially the same as described above. Modifications for microplate format were made as follows: 50 μl samples were incubated overnight at 37°C in either 96-well plate or individual Eppendorf 1.5 ml tubes in demethylation buffer plus peptide, histones or purified nucleosomes. Samples were then diluted to 200 μl with 150 μl of PBS. The samples were then crosslinked to a maleic anhydride amine binding plates (Pierce) in duplicates and prepared following the vendor’s protocols. Plates were then treated for standard ELISA assay by blocking in PBS, 0.1% Tween20 and 2% BSA for 1 hr at room temperature. Antibodies, either methyl specific or pan-H3 (loading control), were then added to separate wells. Antibody to Rabbit IgG conjugated to either Alexa 488 or Alexa 647 were used at a 1:10000 dilution. Relative fluorescence intensities were recorded using a BMG Omega Fluorstar plate reader. For quantification, relative fluorescence units (RFUs) were standardized using a protein concentration standard curve (5, 2, 1, 0.1, 0.01, 0.001, 0.0001, and 0.00001 mg of either histones or nucleosomes). Additionally, two blanks were used to set the non-specific background: demethylation buffer alone and KDM5b alone (no histone added). The negative control was either bulk histones or nucleosomes were incubated in buffer alone. These samples are then normalized to pan-H3. The pan-H3 normalized number is then compared between experimental and the histone/nucleosome alone control. All numbers are expressed as fold change vs. histone/nucleosome only sample. All p-values represent students t-test of experimental vs. control (nucleosomes alone or histone alone).

##### Identification of H2BK43me2/3 and H3K4me2/3 in cells

For *in vivo* identification of H2BK43 and H3K4 methylated peptides, nucleosomes or histones were purified from mouse ES cells (mESCs). For H2BK43me2/3 analysis the nucleosomes were digested with GluC in 1M Guanidine HCl. This yielded the SYSIYVYKVLKQVHPD peptide which was then detected by MRM-MS by monitoring 54 precursor-to-product ion transitions (Table S4). For H3K4 analysis the nucleosomes were digested with ArgC in 0.1M GuHCl yielding the TKQTAR peptide. The peptide was then detected by MRM-MS by monitoring 31 precursor–to-product-ion transitions (Table S5). MRM-MS peak areas were used to compare the relative abundance of H2BK43me2 vs. H2BK43me3 and H3K4me2 vs. H3K4me3 in samples prepared from mESCs at different days of differentiation.

##### Western blotting

For Western blotting, antibodies against KDM5b (at 1:3000 dilution; Bethyl laboratories, cat# A301-813A), H2BK43me2 (at 1:500 dilution; YenZym antibodies), H2B (at 1:1000 dilution; EMD Millipore, cat# 07-371), H3K27me2 (at 1:1000 dilution; Abcam, cat# ab24684), H3K4me3 (at 1:1000 dilution; EMD Millipore, cat# 04-745) and H3 (at 1:1000 dilution; EMD Millipore, cat# 07-690) were used. An HRP-conjugated secondary antibody (BioRad) against either mouse or rabbit IgG was used at 1:10000 dilution. Western blots were detected by ECL (GE Healthcare).

##### Bacterial protein Expression and Purification

Mouse KDM5b was cloned into pDEST17 using the Invitrogen’s Gateway cloning system as previously described (Dey et al., 2008). Recombinant protein was expressed in *E. coli* and purified using Qiagen Ni^+2^ agarose as per vendor protocol (Qiagen). Recombinant proteins were dialyzed in demethylation buffer overnight at 4°C prior to use in enzymatic assays.

##### Statistical Analysis

Statistical analyses were done using Microsoft excel. All p-values were derived from student’s t-tests using two-sample, equal variance, two-tailed settings.

