## Supplemental material for "Proteome-wide Prediction of Lysine Methylation Reveals Novel Histone Marks and Outlines the Methyllysine Proteome"

### Supplementary Information

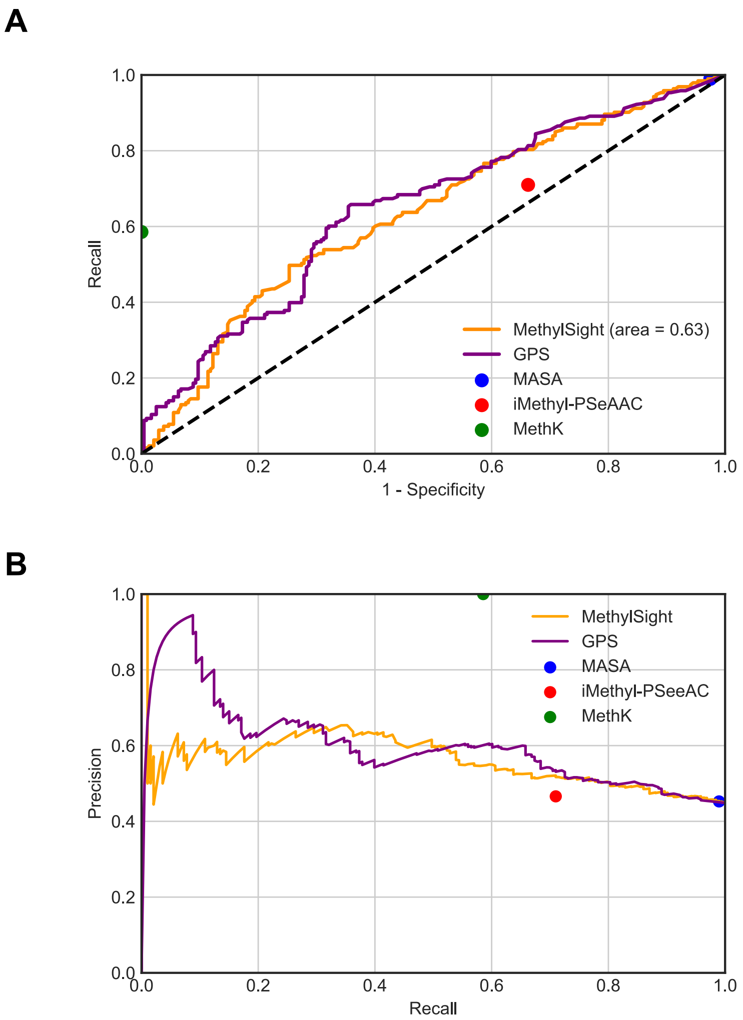


**Fig. S1. Performance curves comparing MethylSight to its competitors over the high-confidence test set.** (A) Receiver operating characteristic (ROC) curves summarizing the anticipated performance of MethylSight and its competitors in terms of recall and specificity. (B) Precision-recall curve illustrating the precision-recall trade-off of MethylSight and other predictors.

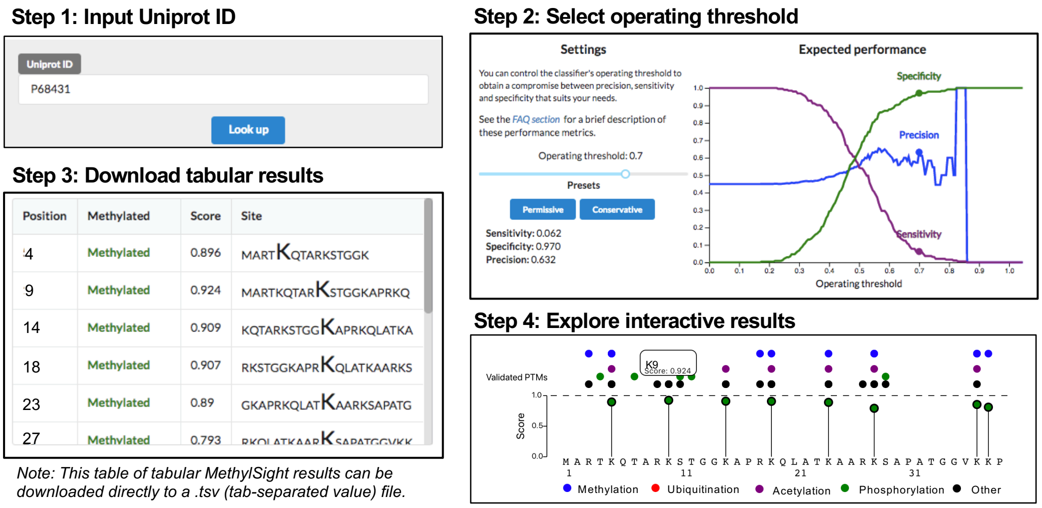


**Fig. S2. Schematic representation of the MethylSight graphical user interface.** MethylSight relies on a user supplied Uniprot identifier to query the prediction (Step 1). Users are then able to select a threshold value based on desired operating performance beyond the preset values for permissive and conservative (0.5 and 0.7, respectively) thresholds (Step 2). Prediction results are then displayed in both a tabular (Step 3) and an interactive (Step 4) format. Histone H3.1 (uniprot P68431) is used here as an example input.

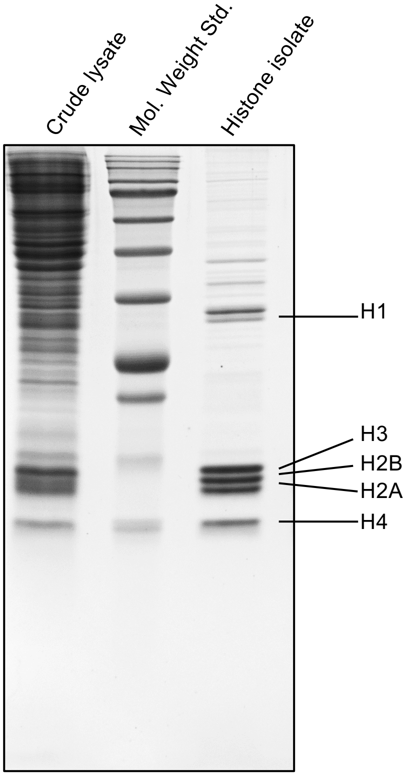


**Fig. S3. Isolation of histones.** Shown is an SDS-PAGE image of the isolated histone proteins (right lane) from the MCF7 cells.

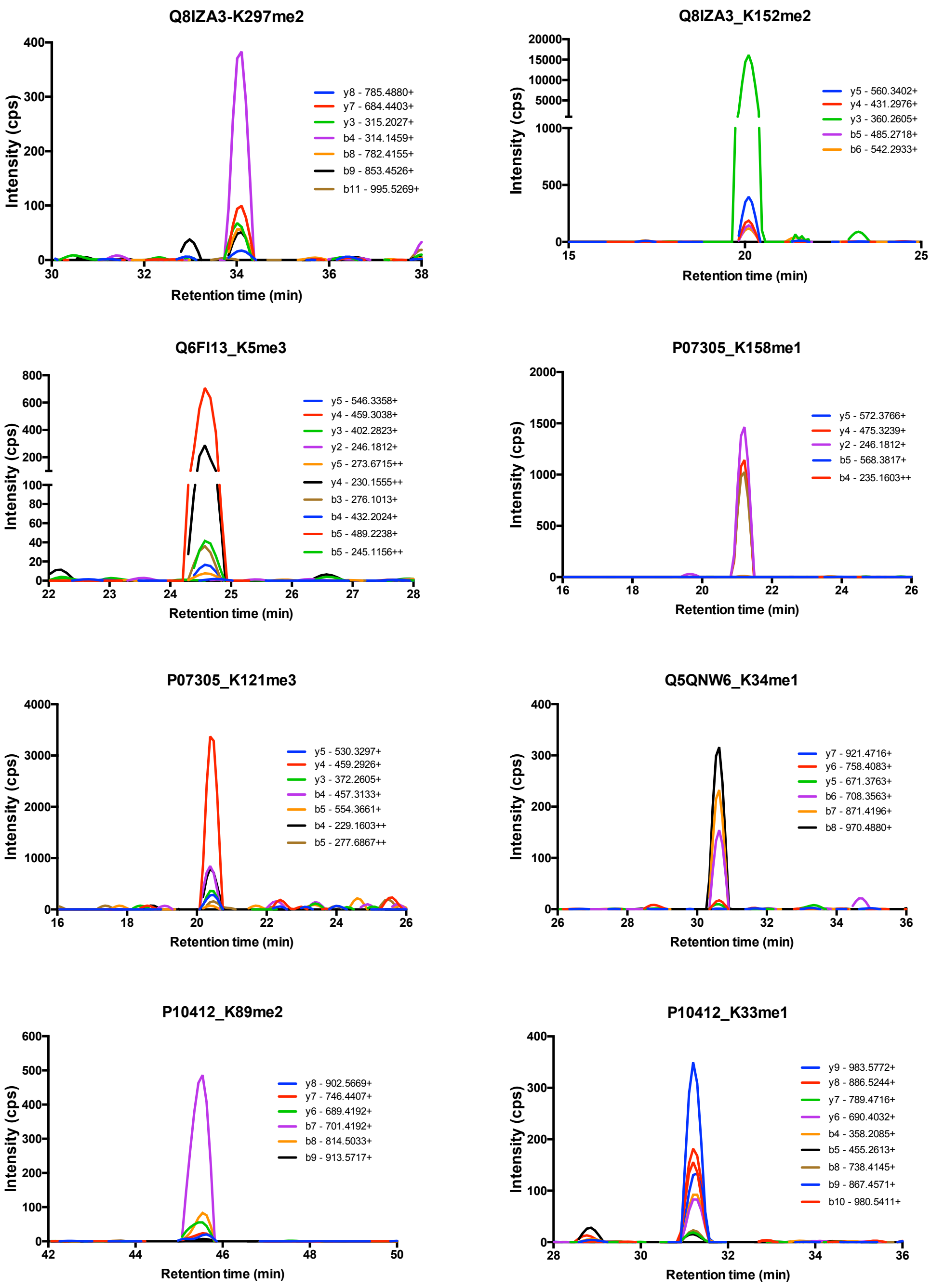


**Fig. S4. Representative MRM transitions used to validate MethylSight predictions of histone H1 Kme sites.** Chromatographs show detected transitions ions that were used to validate indicated histone H1 methylation site. See also Table 2 and Table S2 for further information.

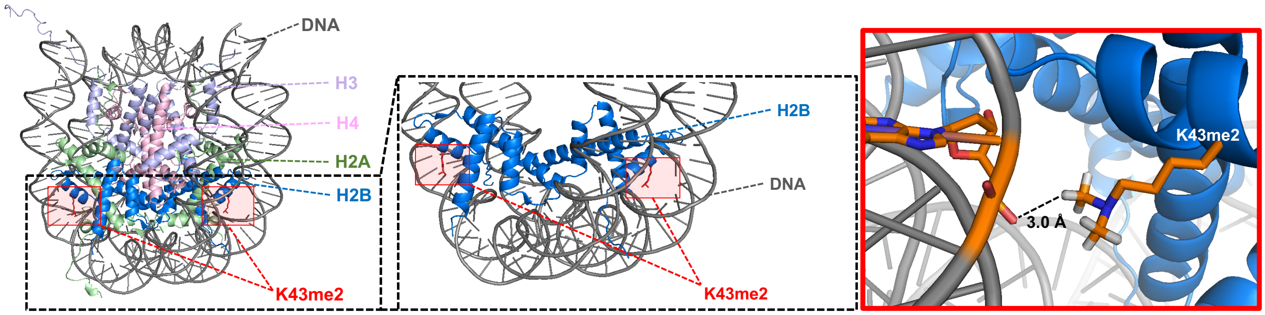


**Fig. S5. H2BK43me2 is in close-proximity to the DNA backbone**. Crystal structure of nucleosome in complex with bound DNA (PDB 1AOI) demonstrating the proximity of the H2BK43me2 methylation site to the DNA phosphate backbone. Interactions modelled with PyMol (v2.2).

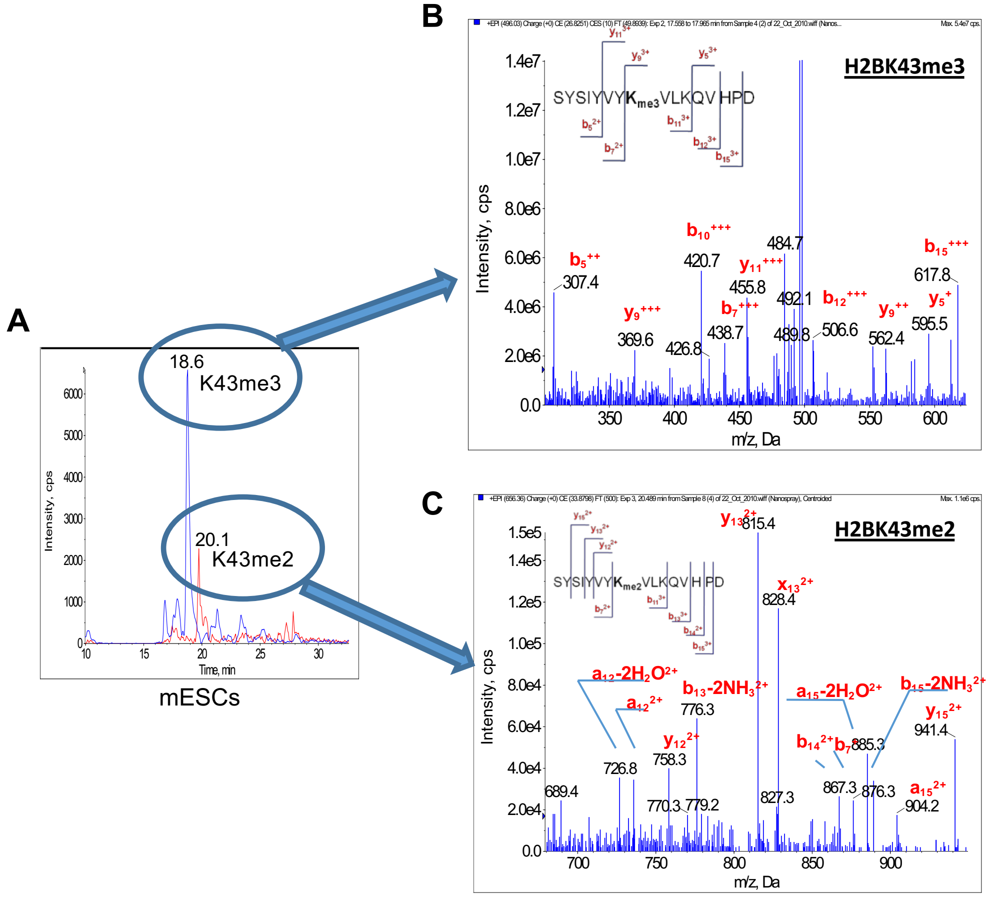


**Figure S6. Confirmation of H2BK43 methylation by MS/MS analysis** **A**. MRM-MS spectra of the H2BK43me3 (blue line) and H2BK43me2 (red line) peptides generated from GluC digestion of nucleosomes purified from mESCs. **B**. MS/MS confirmation of fragments generated for the putative H2BK43me3 peptide. Sequence of the peptide (corresponding to residues 36-52 of H2B) detected by MRM and its MS/MS fragmentation pattern are shown. K43 in the peptide is trimethylated. **C**. MS/MS confirmation of fragment ions generated by the putative H2BK43me2 peptide corresponding to residues 36-52 of mouse H2B. K43 is dimethylated.

| Peptide Sequence | | Histone site |
|---|---|---|
| 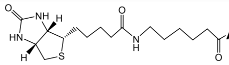 | ART**K**QTARKS | H3K4 |
| 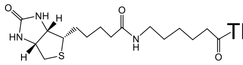 | TKQTAR**K**STGGKA | H3K9 |
| 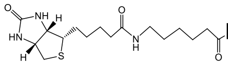 | RKQLAT**K**AARKSA | H3K23 |
| 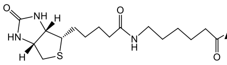 | ATKAAR**K**SAPATG | H3K27 |
| 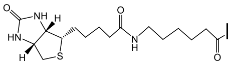 | PATGGV**K**KPHRYR | H3K36 |
| 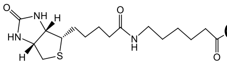 | GAKRHR**K**VLRDNI | H4K20 |
| 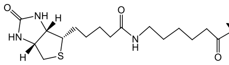 | YVYKVL**K**QVHPDT | H2BK46 |
| 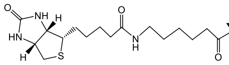 | YVYKVL**K**QVHPDT | H2BK108 |
| 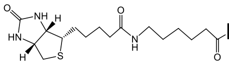 | LPGELA**K**HAVSEG | H2BK43 |

[a]A peptide substrate was derived from the corresponding histone Lys site (identified in red) and synthesized with a biotin moiety coupled to the N-terminus of the peptide with an ε-aminocaproic acid (Ahx) linker. The biotin moiety facilitated peptide isolation prior to mass spec analysis.

**Fig. S7. Sequences of peptide substrates[a] used in the demethylation assay**

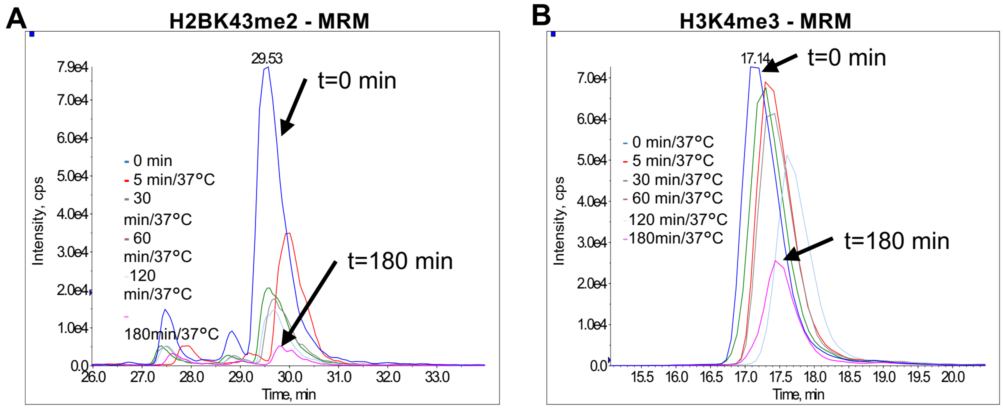


**Fig. S8**. **Mass spectrometric analysis of products from in vitro demethylation assays. (A)** MRM peaks representing precursor-to-product ion transitions (M→ $y_{10}$-$y_4$) of methylated the H2BK43 peptide: Biotin-Ahx-YVY**Kme2**VLKQVHPDT. MRM peaks corresponding to samples taken from the indicated time points of the demethylation assay are shown. (B) MRM peaks representing precursor-to-product ion transitions (M→$y_8$-$y_2$) of methylated H3K4 peptide: Biotin-AHX-ART**Kme3**QTARKS.

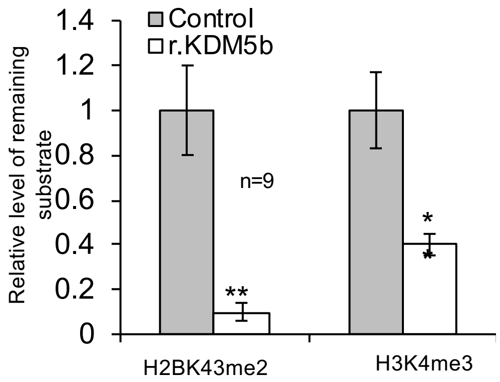


**Fig. S9**. **H2BK43me2 is the preferred substrate for KDM5b in vitro.** Shown are MRM quantification of demethylation of the H2BK43me2 and H3K4me3 peptides mixed at equal molar ratio. **, p<0.01, compared to control; Student's t-test. r.KDM5b, recombinant human KDM5b.

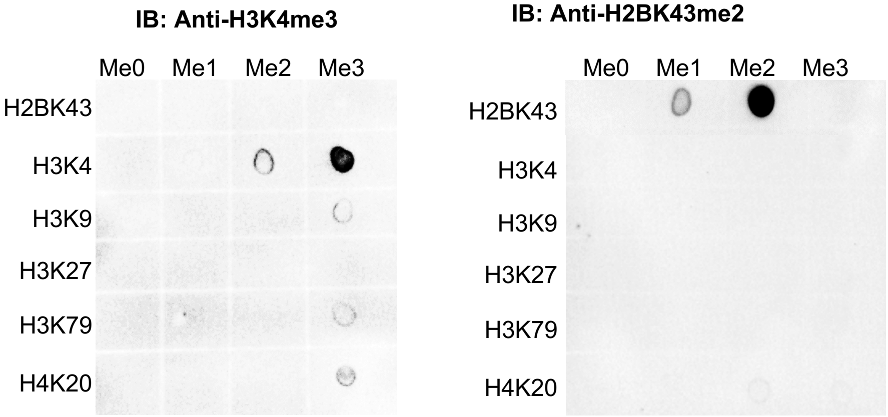


**Figure S10.** ***The anti-H2BK43me2 antibody recognizes the H2BK43me2 mark specifically.*** Histone peptides were spotted on cellulose membrane and probed by anti-H3K4me3 (left) or anti-H2BK43me2 (right) antibody.
